# Phylogenomics defines *Streptofilum* as a novel deep branch of streptophyte algae

**DOI:** 10.1101/2024.03.08.584070

**Authors:** Vojtěch Žárský, Marek Eliáš

## Abstract

Streptophytes constitute a major organismal clade comprised of land plants (embryophytes) and several related green algal lineages. Their seemingly well-studied phylogenetic diversity was recently enriched by the discovery of *Streptofilum capillaum*, a simple filamentous alga forming a novel deep streptophyte lineage in a two-gene phylogeny^1^. A subsequent phylogenetic analysis of plastid genome-encoded proteins resolved *Streptofilum* as a sister group of nearly all known streptophytes, including Klebsormidiophyceae and Phragmoplastophyta (Charophyceae, Coleochaetophyceae, Zygnematophyceae, and embryophytes)^2^. In a stark contrast, another recent report, published in *Current Biology* by Bierenbroodspot et al.^3^, presented a phylogenetic analysis of 845 nuclear loci resolving *S. capillatum* as a member of Klebsormidiophyceae, nested among species of the genus *Interfilum*. Here we demonstrate that the latter result is an artefact stemming from an unrecognized contamination of the transcriptome assembly from *S. capillatum* by sequences from *Interfilum paradoxum*. When genuine *S. capillatum* sequences are employed in the analysis, the position of the alga in the nuclear genes-based tree fully agrees with the plastid genes-based phylogeny. The “intermediate” phylogenetic position of *S. capillatum* predicts it to possess a unique combination of derived and plesiomorphic traits, here exemplified, respectively, by the “Rho of plants” (ROP) signaling system and the cyanobacteria-derived plastidial transfer-messenger ribonucleoprotein complex (tmRNP). Our results underscore *S. capillatum* as a lineage pivotal for the understanding of the evolutionary genesis of streptophyte, and ultimately embryophyte, traits.

---

*Streptofilum capillatum*, described only in 2018 by Mikhailyuk et al.^1^, is a humbly-looking green alga forming short filaments inside a mucilaginous sheath (Figure 1A, inset). A phylogeny built from a combination of the nuclear 18S rRNA and the plastidial *rbcL* genes suggested that *S. capillatum* represents a deep separate streptophyte lineage, even possibly sister to all other streptophytes combined^1^. More recently, Glass et al.^2^ sequenced the plastid genome of *S. capillatum* and corroborated its separate phylogenetic status by analyzing a set of 44 plastid-encoded proteins, placing the alga with full support sister to nearly all other streptophytes combined except for the more basally positioned genera *Mesostigma* and *Chlorokybus*. From this perspective, the results of the phylotranscriptomic analysis reported by Bierenbroodspot et al.^3^ are highly surprising: having *S. capillatum* positioned within Klebsormidiophyceae in a tight clade together with representatives of the genus *Interfilum* (even making the genus paraphyletic), their tree separates the alga from the position implied by the plastid genes-based phylogeny by five fully supported nodes.

**Figure 1.**
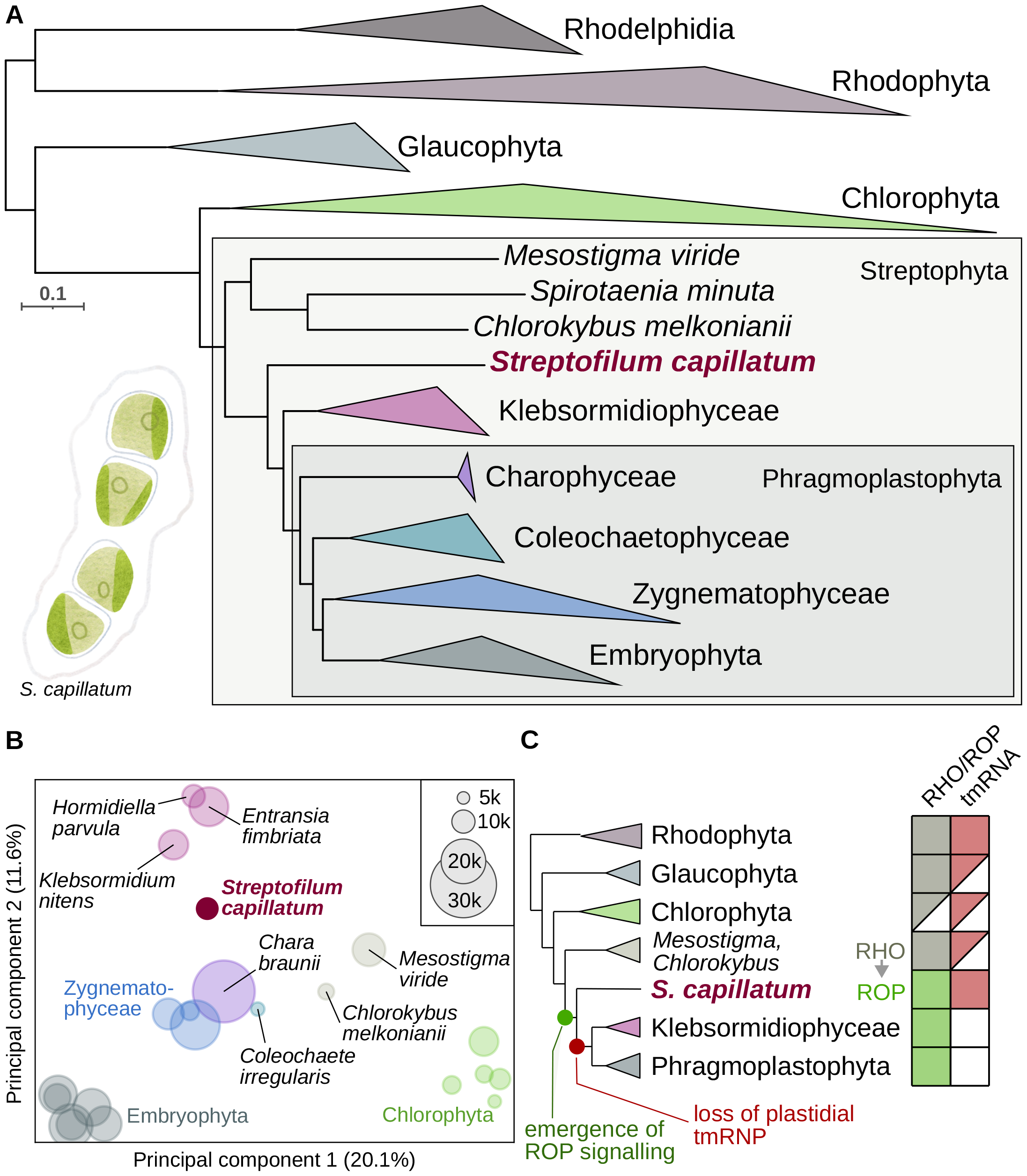
*Streptofilum capillatum* represents a unique, deeply diverged streptophyte lineage. (A) A phylogeny of 184 archaeplastid taxa based on a supermatrix built from 96 conserved nucleus-encoded proteins (24,431 amino acid positions). The tree was inferred using the maximum likelihood method and the LG+C60+G mixture model. Branch support was assessed with ultrafast bootstrap approximation (UFBoot), with nearly all branches, including all explicitly shown in the figure, having received maximal support (UFBoot value of 100). The scale bar indicates the number of substitutions per site. For simplicity, clades outside the focus of the study were collapsed and are displayed as triangles; a full version of the tree is provided as Figure S2. Sequences from the decontaminated *S. capillatum* transcriptome assembly form a distinct lineage within streptophytes sister to Klebsormidiophyceae and Phramoplastophyta combined (see Figure S2 for a detailed view of the phylogenetic relationships within Klebsormidiophyceae, including the position of the branch corresponding to the klebsormidiophycean contaminant in the original *S. capillatum* transcriptome assembly). The inset shows a drawing of *S. capillatum* (a short filament consisting of cell dyads in a common mucilaginous covering); the figure was drawn based on original photos published by Mikhailyuk et al.^1^. (B) Comparison of protein repertoires (predicted proteomes) encoded by selected green algae and plants. The organisms were plotted according to the principal component analysis of a matrix representing the sharing of orthologous groups among the species. The size of the circles indicates the size of the non-redundant proteome (considering only protein sequences longer than 200 amino acid residues) predicted for the organism (see the scale in the top right corner). A more detailed version of the plot is provided as Figure S5. (C) Two exemplar traits exhibiting a unique combination of states in *S. capillatum*. The states are mapped onto a schematic phylogeny that reflects the results shown in panel A. The squares to the right of the tree show the distribution of two forms of Rho GTPase signaling (Rho – the ancestral one; ROP – the derived “Rho of plants” type) and of the plastidial tmRNP complex in the respective taxa. Presence/absence is indicated by filled/empty squares, respectively, half-filled squares means that the given trait is present only in some representatives of the taxon.

We hypothesized that the result by Bierenbroodspot and colleagues may be an artefact, which was indeed corroborated by our scrutiny (detailed in Supplementary Methods) of the transcriptome assembly generated by the authors for *S. capillatum* (strain SAG 2559). Specifically, we found the assembly to be contaminated by sequences from *Interfilum paradoxum*, most likely the SAG 338-1 strain that was subjected to transcriptome sequencing in the same study. This is documented by the presence of two different 18S rRNA sequences in the data, one corresponding to *S. capillatum* and the other identical to the *I. paradoxum* sequence (Figure S1). Analyses of the predicted protein-coding genes also revealed the presence of a large number of pairs of homologs, one with a lower read coverage and exhibiting identity or extreme similarity to sequences from *I. paradoxum* or other reference klebsormidiophyceans, and the other with a higher coverage and being less similar to klebsormidiophycean sequences yet still clearly of a green algal provenance, thus apparently coming from *S. capillatum* itself. As we found out, the phylogenomic dataset employed by Bierenbroodspot and colleagues is dominated by contaminant-derived sequences, explaining both the position and the length of the *S. capillatum* branch in their tree.

We bioinformatically separated the *S. capillatum* and *I. paradoxum* components of the mixed assembly, added orthologs from both sequence subsets to a supermatrix of 96 nucleus-encoded proteins, and inferred a tree from it (Figure 1A, Figure S2). Most importantly, the lineage corresponding to putative bona fide *S. capillatum* sequences was found (with full statistical support) exactly where the alga is expected to sit based on the plastid genome-based phylogeny. Our analyses thus corroborate *S. capillatum* as a novel separate streptophyte lineage sister to Klebsormidiophyceae and Phragmoplastophyta combined. The most recent estimate places the split between Klebsormidiophyceae and Phragmoplastophyta to ∼896 mya^4^, so the *S. capillatum* lineage must have diverged from other extant streptophytes even earlier, yet has not diversified or has experienced extinction, surviving in a single presently known species. However, further work may ultimately unravel its additional representatives, as *S. capillatum* itself had been overlooked for so long despite occurring in a mundane habitat (the original strain was isolated from arable field soil in the Czech Republic^1^).

A comparison of encoded protein repertoires supports the closer relationship of *S. capillatum* to Klebsormidiophyceae and Phragmoplastophyta than to other green algae (Figure 1B). *S. capillatum* thus likely exhibits many genetic traits so far considered to be evolutionary innovations of its sister group, but its phylogenetic position at the same time implies it may have retained plesiomorphies lost in its sister lineage. Here we provide one example of each trait type (Figure 1C). A recent study pinpointed a major change in the Rho GTPase-based signalling machinery, critical for cell morphogenesis, to have happened in early streptophyte evolution after the split of the *Mesostigma*/*Chlorokybus* lineage^5^. We show that *S. capillatum* shares with Klebsormidiophyceae and Phragmoplastophyta both key attributes (Figure S3), i.e. ROP GTPase that has evolved by transformation of the ancestral streptophyte Rho protein, and a novel ROP-specific GTPase regulator (RopGAP). As the exemplar *S. capillatum* plesiomorphy we report the identification of an *ssrA* gene in its plastome missed in the original annotation^2^ (Figure S4) and of the functionally associated nucleus-encoded SmpB protein (Figure S5). Like in bacteria, the *ssrA* product (tmRNA) and SmpB (as parts of the tmRNP complex) mediate rescue of stalled plastid ribosomes^6^. The *ssrA* gene occurs in plastomes of various other algae including *Mesostigma*, but not Klebsormidiophyceae and Phragmoplastophyta^7,8^, and SmpB follows this distribution. A future systematic analysis, ideally based on a complete *S. capillatum* genome, will certainly further refine the reconstructions of the trait evolution in streptophytes.

## ACKNOWLEDGEMENTS

We thank Alina Žárská for the *S. capillatum* drawing. This work was supported by the Czech Science Foundation (project 23-05764S) and the European Union under the LERCO project number CZ.10.03.01/00/22_003/0000003 via the Operational Programme Just Transition.

