## Supplemental material for "Phylogenomics defines *Streptofilum* as a novel deep branch of streptophyte algae"

**This file contains Supplementary Methods, Supplementary References, and Supplementary Figures. Supplementary Tables are provided in a separate xlsx file.**

### Supplementary Methods

*18S rRNA analysis*

Searching the original transcriptome assembly from *Streptofilum capillatum* SAG 2559 (Bierenbroodspot et al. 2024) with the previously reported 18S rRNA gene sequence from this strain (MG652626.1) as a BLASTN (Altschul et al. 1990) query identified multiple contigs corresponding to the 18S rRNA gene, which were revealed by further scrutiny to be derived from various eukaryotic species including *S. capillatum* itself and to represent various putatively chimeric combinations of regions derived from different species. We therefore reassembled the original RNA-seq reads (NCBI SRA, SRR26030666) with rnaSPAdes (Bushmanova et al. 2019) using the "-k 21,33,55,77,99,127" setting (while the original assembly was obtained by employing Trinity). Ribosomal RNA sequences were then detected in the new assembly by searching the transcripts against the covariance models representing different rRNA families from the Rfam database (Kalvari et al. 2021) using Infernal cmscan (Nawrocki and Eddy 2013; https://www.ebi.ac.uk/jdispatcher/rna/infernal_cmscan). The rRNA sequences were then annotated by searching against the NCBI nt database using BLASTN. The analysis revealed the presence of contigs corresponding to 18S rRNA sequences from *Streptofilum capillatum* (read coverage 11892×; 100% identity to MG652626.1), the klebsormidiophycean *Interfilum paradoxum* (coverage 1194×; 99.3% identity to EU434016.1), the oomycete *Globisporangium* sp. (coverage 1528×; 99.9% identity to EF023430.1), the amoebozoan *Acanthamoeba* sp. (coverage 174×; 98.9% identity to AY173015.1), and the fungus *Sarcocladium* sp. (coverage 43×; 99.9% identity to LFCJ01011078.1), indicating a complex taxonomic composition of the transcriptome data attributed to *S. caillatum*. A phylogenetic tree was constructed from a set of 18S rRNA sequences from the *S. capillatum*, the putative *I. paradoxum* contamination, and a selection of plant and green algal species. The sequences were aligned using MAFFT (Katoh and Standley 2013), the alignment was trimmed using trimAl (Capella-Gutiérrez et al. 2009) with the "-automated1" option, and a ML phylogenetic tree was built using IQ-TREE 2 (Minh et al. 2020) using the GTR+F+I+G4 model. The phylogenetic tree was visualized and edited in iTOL (Letunic and Bork 2021); this program was used to also process all other trees presented in this study.

*Phylogenomic analysis*

Protein sequences derived from the *S. capillatum* SAG 2559 transcriptome assembly, generated and initially decontaminated by Bierenbroodspot et al. (although failing to account for the *I. paradoxum* contamination) were searched against the predicted proteomes from *Klebsormidium nitens* NIES-2285 and *Interfilum paradoxum* 338-1 using Diamond BLASTP (Buchfink et al. 2021). The proteins were then separated into two sets: sequences with a high identity to Klebsormidiophyceae reference proteomes (>90% identity), assumed to correspond to *I. paradoxum* contaminants, and sequences with a lower (<80%) identity to the Klebsormidiophyceae reference proteomes presumably corresponding to genes from *S. capillatum*, with possibly residual contaminants from other eukaryotes (see above). The former set was further denoted as "*Interfilum* 2559" in the phylogenomic analysis, while the latter as "*Streptofilum capillatum*" (Table S1A).

A recent 96-gene phylogenomic dataset employed for reconstructing the phylogeny of Archaeplastida (Bowles et al. 2024) was used as a starting point for our phylogenomic analysis. Profile HMMs, built for the alignments of each of the 96 sets of orthologous proteins, were used to search with HMMER (Eddy 2011) against sets of predicted proteomes from additional organisms (listed in Table S1A) including *S. capillatum* and the *I. paradoxum* contaminant (“*Interfilum* 2559”). The best hit, if having e-value <1e-50, was selected for each of the new organism and added to the phylogenomic dataset, assuming that green algal sequences will be preferentially selected over possible contaminating homologs from non-Chloroplastida eukaryotes. Each set of homologs was then aligned with MAFFT, trimmed with trimAl and a phylogeny was inferred for it with FastTree 2 (Price et al. 2010). The single-gene phylogenies were manually checked to identify obvious contaminants, paralogs, and outliers, which were then removed. 55 genes with high confidence assigned to *S. capillatum* and 44 genes with high confidence assigned to the *I. paradoxum* contaminant in the SAG 2559 transcriptome assembly were identified and added to the final phylogenomic dataset (see Table S1A for additional information on the phylogenomic dataset). Sequences of each of the orthologous groups were filtered for non-homologous characters with PREQUAL (Whelan et al. 2018) and aligned with MAFFT using the “--localpair --maxiterate 1000” options. The alignments were then filtered with Divvier (Ali et al. 2019) to identify clusters of high confidence homologies and finally trimmed with trimAl using the “-automated1” setting. A phylogenomic supermatrix was built by concatenating these highly refined alignments and a phylogenetic tree was inferred with IQ-TREE 2 using the LG+C60+G mixture model (Quang et al. 2008) and the branch support was assessed with ultrafast bootstrap approximation (UFBoot) (Hoang et al. 2018). The complete phylogenomics dataset and single-gene phylogenies can be downloaded from Figshare (https://figshare.com/s/1b1a1672c26a96f10ee8).

*Transcriptome decontamination*
A new decontaminated dataset of *S. capillatum* transcripts and predicted proteins was prepared based on our rnaSPAdes re-assembly of the sequencing reads generated by Bierenbroodspot et al. (see the 18S rRNA analysis section for details on the assembly). The coding regions within transcripts were predicted using TransDecoder (Haas BJ, https://github.com/TransDecoder/TransDecoder) with hints provided by homology searches against the UniProt reference proteomes (UniProt Consortium 2019) and the Pfam database of protein families.

All contigs were searched with BLASTN against a selection of reference genome or transcriptome assemblies from the suspected contaminants or their relatives: two Klebsormidiophyceae reference assemblies (the genome of *Klebsormidium nitens* - GCA_000708835 and the transcriptome of *Interfilum paradoxum* SAG 338-1 generated by Bierenbroodspot et al.), two *Acanthamoeba* genomes (*Acanthamoeba castellanii* - GCA_021020595 and *A. polyphaga* - GCA_001567625), two *Globisporangium* genomes (*Globisporangium emineosum* - GCA_023334195 and *G. nagaii* - GCA_023335995), and two *Sarocladium* genomes (*Sarocladium kiliense* - GCA_030734395 and *S. strictum* - GCA_030435815) using BLASTN. Additionally, all contigs were searched against two Phragmoplastophyta genomes (*Spirogloea muscicola* - GCA_009602725 and *Physcomitrium patens* - GCA_000002425). Contigs were also searched against UniProt Reference proteomes, EukProt (Richter et al. 2022), and a selection of predicted proteomes from Chloroplastida representatives (listed in Table

S1B, excluding *S. capillatum*) with Diamond BLASTX. Finally, TransDecoder-predicted protein sequences were searched against UniProt Reference proteomes, EukProt, and the selection of Chloroplastida proteomes with Diamond BLASTP.

Of the 105,416 contigs in the full assembly, 511 were removed due to having a ribosomal RNA gene that was not previously annotated as representing *S. capillatum*. 63,400 contigs were removed due to having a significant stretch (>50% of contig length) covered with significant (e-value <1e-20) high-identity (>70% sequence identity) BLASTN hits against either of the contaminant reference assemblies while having low identity to the Phragmoplastophyta reference assemblies. 3,968 contigs were removed because their best BLASTX hit was outside of Chloroplastida and was significant (e-value <1e-10) with high identity (>80%). All significant (e-value <1e-10) best BLASTX hits derived from Centramoebida, Oomycota, or Pezizomycotina species were also removed in this step. 9,368 contigs were removed because their length was smaller than 300 bps. This decontamination process resulted in a set of 28,169 contigs. Of the 45,223 protein sequences predicted by TransDecoder based on the selected contigs, 14,979 were removed because they were shorter than 100 amino acids while having no significant BLASTP hit (e-value >1e-5), resulting in a set of 30,244 putative proteins considered to represent the predicted *S. capillatum* proteome. The filtered dataset can be downloaded from Figshare (https://figshare.com/s/1b1a1672c26a96f10ee8).

*Whole-proteome comparisons*
Orthologous groups were reconstructed *de novo* for a set predicted proteomes from selected Chloroplastida representatives, including the decontaminated transcriptome assembly from *S. capillatum* (Table S1B) using Broccoli v1.2.1 using "-sp_overlap 1" (Derelle et al. 2020). The occurrences of orthologous groups in each proteome were coded as a binary matrix, which was then subjected to a Principal component analysis using the Scikit-learn Python library (Pedregosa et al. 2011). The organisms were then plotted in 2D based on the first two principal components (Figure 1B, Figure S6). The size of the organism circles in Figure 1B and Figure S6 correspond to the size of the filtered non-redundant proteome, which was acquired by filtering sequences at 90% identity with cd-hit (Fu et al. 2012) and removing sequences with length bellow 200 amino acids. The protein counts, filtered protein counts, and counts of orthologous groups recovered by Broccoli for each proteome can be found in Table S1B. Counts of shared Broccoli orthologous groups between proteomes can be found in Table S1C.

*Phylogenetic analyses of Rho GTPases, RhoGAP proteins and SmpB*
Rho GTPase homologs were searched with TBLASTN in the transcriptome assemblies from *S. capillatum* (the decontaminated version) and *I. paradoxum*, revealing a single gene in each species. For phylogenetic analysis, the respective protein sequences were combined with the set of Rho GTPase sequenced analyzed by Mulvey and Dolan (2023). Sequences with the RhoGAP domain were identified in the predicted proteomes from *S. capillatum*, *I. paradoxum*, and *Spirotaenia minuta* using BLAST and HMMER searches with a profile HMM built from the seed alignment of the Pfam family PF00620 (https://www.ebi.ac.uk/interpro/entry/pfam/PF00620/entry_alignments/). For *I. paradoxum* the transcriptome assemblies generated by both Bierenbroodspot et al. (2024) and the OneKP project (One Thousand Plant Transcriptomes Initiative 2019) were analyzed and a representative sequence (more complete or without retained introns) for each gene was selected from one or the other assembly. For phylogenetic analysis, the sequences were combined with a set of RhoGAP domain-containing protein sequences analyzed by Mulvey and Dolan (2023) and an extra sequence from *Klebsormidium nitens* omitted by the authors by identified by us a orthologous to some of the other streptophyte proteins included in their dataset. BLAST was used to identify candidate plastidial SmpB genes in genome and transcriptome assemblies from Chloroplastida members in the NCBI databases, in transcriptome

assemblies generated by the OneKP project, and in the transcriptome assemblies reported by Bierenbroodspot et al. (2024). In a parallel, a profile HMM built from the seed alignment of the Pfam family PF01668 (=SmpB; https://www.ebi.ac.uk/interpro/entry/pfam/PF01668/) was used for a HMMER search against the EukProt and hits above the inclusion threshold were retrieved. All newly identified SmpB sequences were combined with previously identified eukaryotic SmpB sequences (Gray et al. 2020) and a selection of bacterial homologs retrieved as the best 100 hits in BLASTP searches with the plastidial SmpB from *S. capillatum* and the mitochondrial SmpB from *Ancoracysta twista* as queries against the nr_bac70 protein sequence database (a version of the NCBI non-redundant sequence database filtered for bacterial sequences and a maximum pairwise sequence identity of 70%) at https://toolkit.tuebingen.mpg.de/tools/psiblast. A preliminary phylogenetic analysis using the "Phylogenetic analysis pipeline by ETE3" (with FastTree as the tree inference method; https://www.genome.jp/tools-bin/ete) was performed to identify and remove bacterial contaminants in the eukaryotic datasets and to select representative plastidial SmpB proteins from taxa other than Chloroplastida. Collected sets of Rho GTPase and SmpB sequences were aligned with MAFFT using the "--maxiterate 1000 --localpair" option, the alignment was trimmed with Trimal using the "-automated1" option, whereas hmmalign (from the HMMER package) with the RhoGAP profile HMM was applied (with the --trim option) to obtain a multiple alignment of the RhoGAP domains in the proteins (while trimming the non-homologous regions of the sequences). ML phylogenetic trees were inferred from the alignments with IQ-TREE 2 using the LG+I+G4 model and 10000 UFBoot replicates.

*Identification of the plastidial* ssrA *gene*
The expanded list of plastidial SmpB homologs gathered by our analyses suggested that an *ssrA* gene resides in plastomes of several taxa where it was previously not identified, including *Tetraselmis* spp. in the class Chlorodendrophyceae (Turmel and Lemieux 2018) and *S. capillatum* (Glass et al. 2023). In addition, further SmpB homologs pointed to the existence of plastidial *ssrA* in taxa where the plastome sequence was previously not analysed (*Protoeuglena noctilucae* and *Picocystis* sp. ML) or is unavailable (*Scourfieldia* sp.). The plastomes suspected to contain an *ssrA* gene were thus subjected to a search against the Rfam database using Infernal cmscan, which identified a highly significant hit (e.g., e-value of 2.3e-26 for *S. capillatum*) to the tmRNA model in all cases, except for *Tetraselmis* spp. However, regions obviously fitting the sequence pattern of the *ssrA* gene were identified in all four *Tetraselmis* plastome sequences available for analysis (see Figure S4) by checking the putative intergenic region downstream of the *rbcL* gene, i.e. where *ssrA* normally resides in Chlorophyta plastomes (Turmel et al. 2015), indicating that the current tmRNA covariance model in Rfam is not sensitive enough to detect more divergent organellar *ssrA* genes. Sequences of all the (putative) *ssrA* genes from green algal plastomes were aligned with MAFFT and for display the alignment was annotated following the scheme in the supplementary Figure S4 in Turmel et al. (2015).

### Supplementary figures

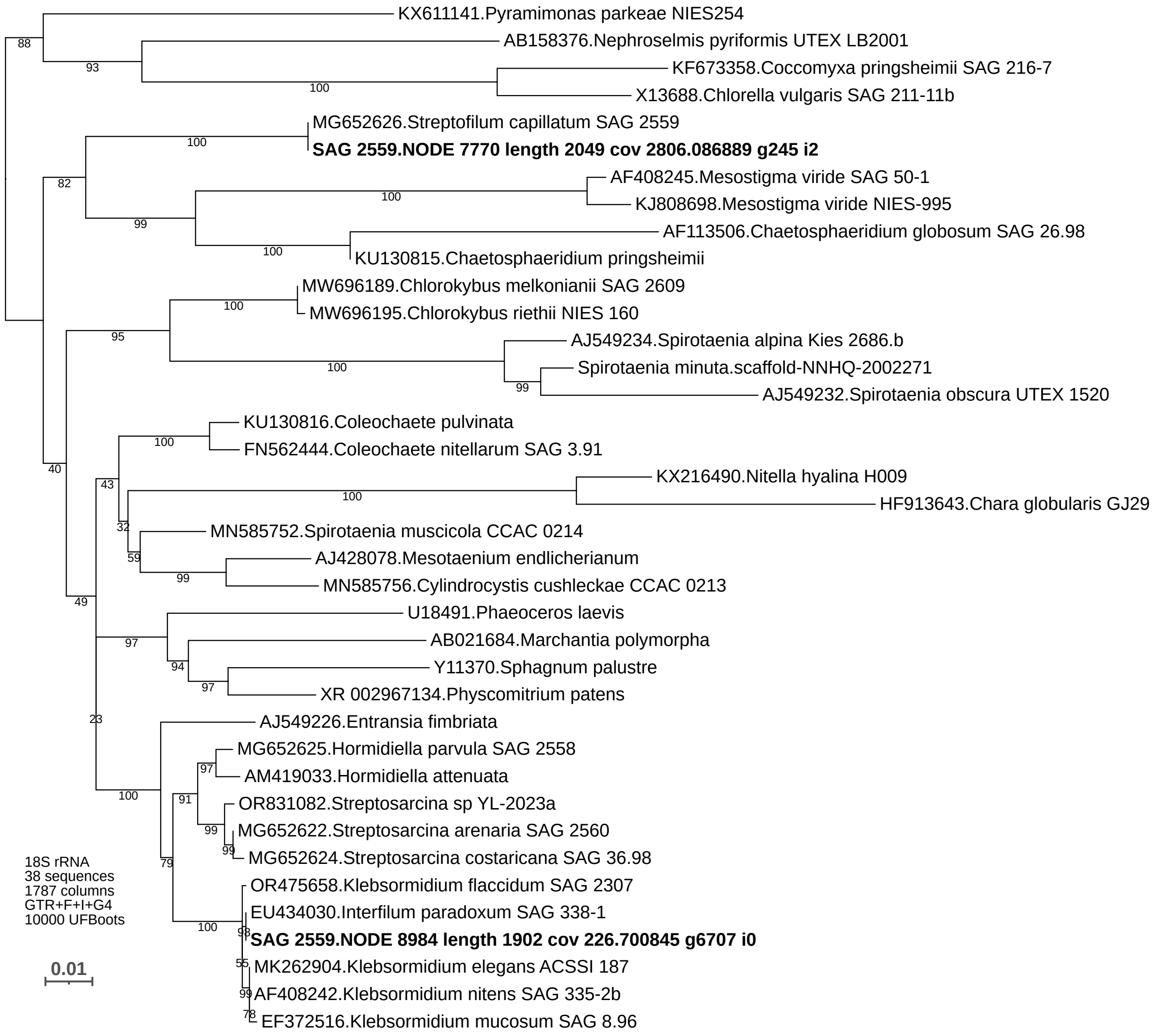


**Figure S1. Maximum likelihood phylogeny inferred from sequences of the 18S rRNA gene from selected Chloroplastida representatives.** Five reference sequences from Chlorophyta were included to provide an outgroup for a more systematically sampled phylogenetic diversity of Streptophyta. The two sequences highlighted in bold correspond to two different green algal 18S rRNA sequences identified in our reassembly of the original RNA-seq reads reported from *Streptofilum capillatum* SAG 2559 (the sequence IDs indicate the respective contigs in the assembly). One of these sequences is identical to the 18S rRNA sequence reported from *S. capillatum* in the original report on the species, whereas the other is identical to the 18S rRNA gene from *Interfilum paradoxum*. Note the difference in the read coverage of the two contigs, consistent with the notion that the less abundantly represented *I. paradoxum*-like sequence comes from a contaminant.

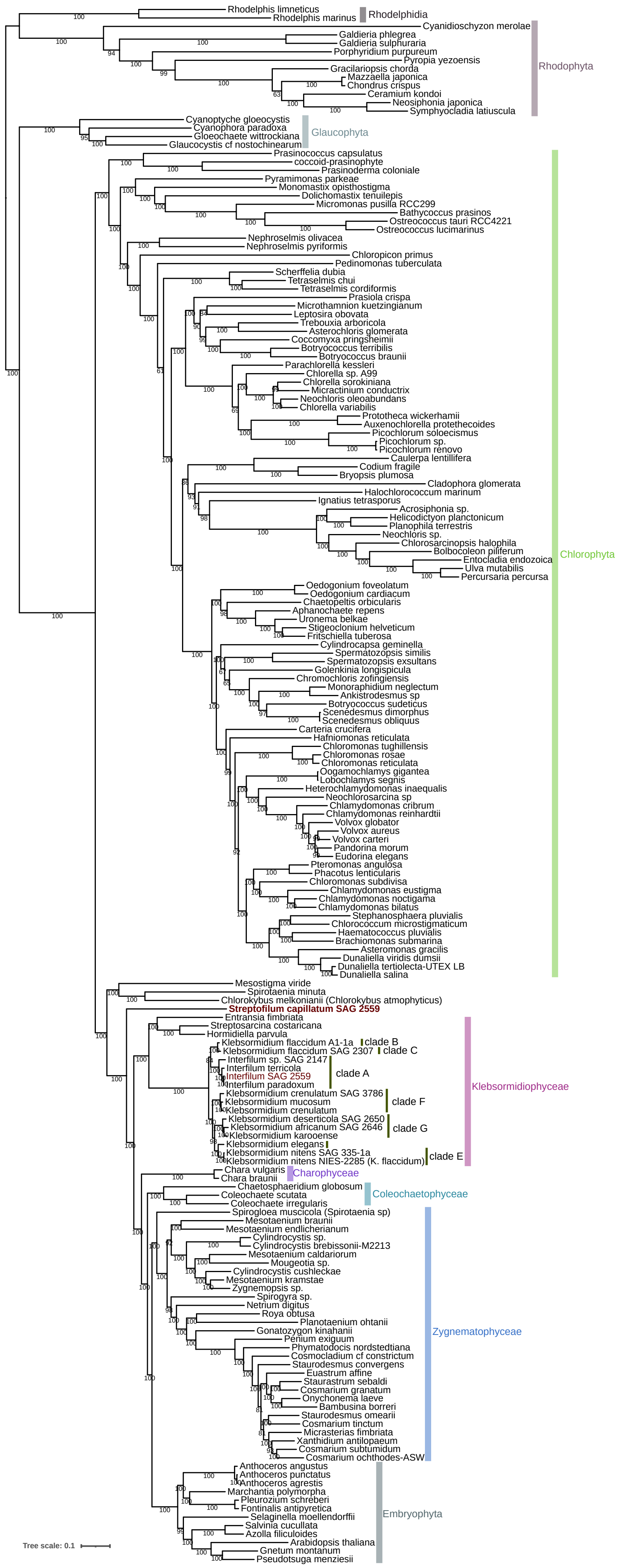


**Figure S2. Maximum likelihood phylogeny of Archaeplastida inferred from a supermatrix built from 96 conserved nucleus-encoded proteins.** Displayed is a full version of the phylogenomic tree presented as Figure 1A in the main text (see the legend to that figure for further details on the tree inference and display conventions). Note the identity of the "*Interfilum*" component contaminating the originally published transcriptome assembly from *Streptofilum capillatum* SAG 2559 with *Interfilum paradoxum* SAG 338-1 (in Klebsormidiales "clade A").

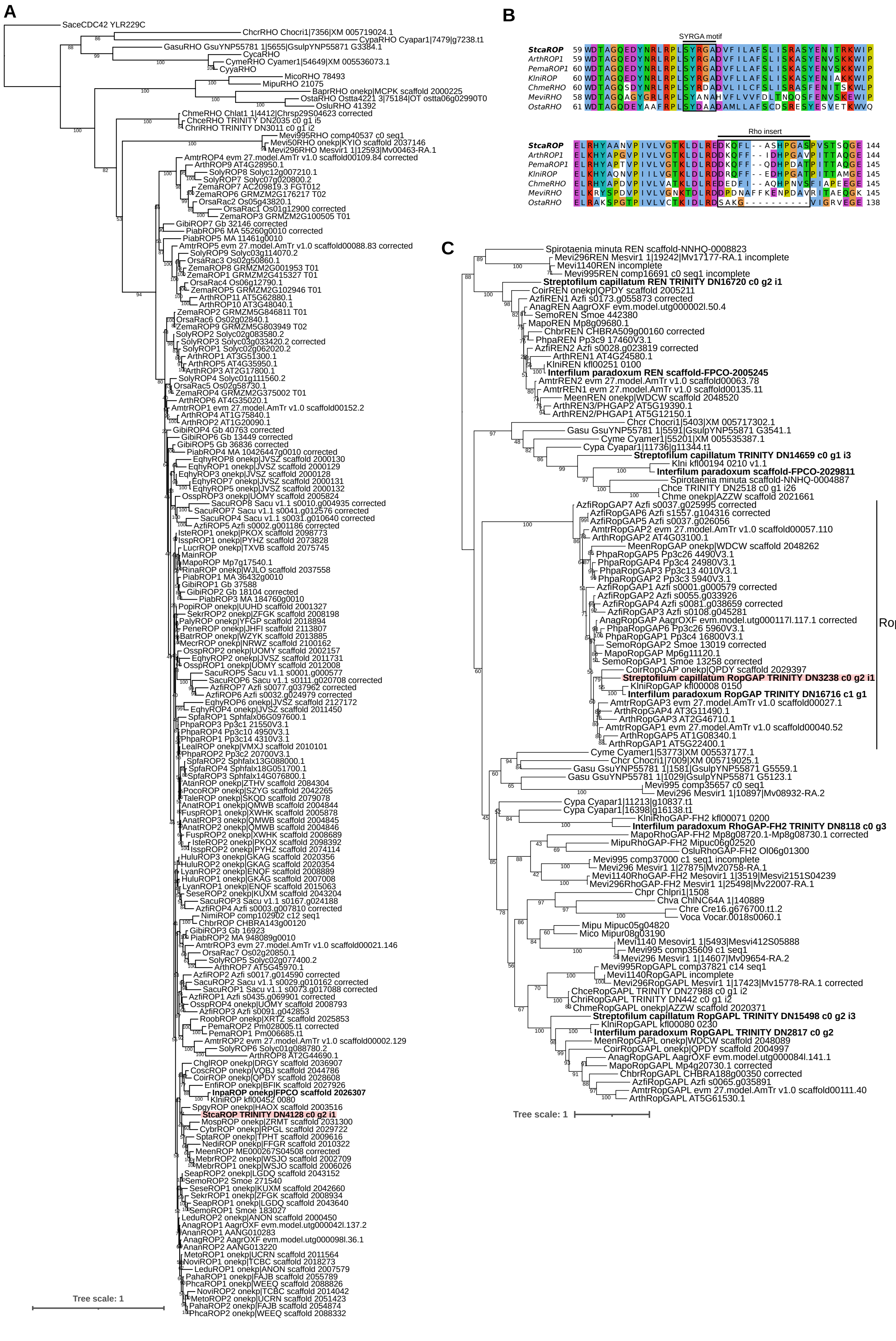
A
B
C
SYRGA motif
Rho insert
RopGAP
Tree scale: 1
Tree scale: 1

**Figure S3. The presence of ROP signalling in *Streptofilum capillatum* SAG 2559.** (A) Maximum likelihood phylogeny of Rho family GTPases demonstrating that the single Rho GTPase found in the decontaminated *S. capillatum* transcriptome assembly (highlighted in red) is affiliated with ROP-type GTPases from other streptophytes. Note that the sequence differs from ROP sequences found in Klebsormidiophyceae (including *Interfilum paradoxum*; InpaROP), ruling out the possibility that it represents an unrecognized contamination in the transcriptome assembly. (B) The *S. capillatum* Rho family GTPase exhibits the SYRGA motif and the 12-aa-long Rho insert, signatures of ROP GTPases. (C) Maximum likelihood phylogeny of RhoGAP domain-containing proteins demonstrating that *S. capillatum* encodes the ROP-specific regulator RopGAP (highlighted in red). Again, a different sequence was identified in the transcriptome assembly from *I. paradoxum*, supporting the authenticity of the sequence assigned to *S. capillatum*. All sequences from *S. capillatum* and *I. paradoxum* are typed in bold. The four-letter abbreviations for sequences in A-C are as follows: *Amborella trichopoda* (Amtr), *Anthoceros agrestis* (Anag), *Arabidopsis thaliana* (Arth), *Azolla filiculoides* (Azfi), *Chara braunii* S276 (Chbr), *Chlorokybus cerffii* SAG34.98 (Chce), *Chondrus crispus* Stackhouse (Chcr), *Chlorokybus melkonianii* CCAC0220 (Chme), *Chloropicon primus* CCMP1205 (Chpr), *Chlamydomonas reinhardtii* (Chre), *Chlorokybus riethii* NIES160 (Chri), *Chlorella variabilis* NC64A (Chva), *Coleochaete irregularis* (Coir), *Cyanidioschyzon merolae* 10D (Cyme), *Cyanophora paradoxa* CCMP329 (Cypa), *Galdieria sulphuraria* YNP5578.1 (Gasu), *Interfilum paradoxum* SAG 338-1 (Inpa), *Klebsormidium nitens* NIES2285 (Klni), *Marchantia polymorpha* (Mapo), *Mesotaenium endlicherianum* SAG12.97 (Meen), *Mesostigma viride* CCAC1140 (Mevi1140), *Mesostigma viride* NIES296 (Mevi296), *Mesostigma viride* NIES995 (Mevi995), *Micromonas commoda* RCC299 (Mico), *Micromonas pusilla* CCMP1545 (Mipu), *Ostreococcus lucimarinus* (Oslu), *Physcomitrium patens* (Phpa), *Saccharomyces cerevisiae* (Sace), *Selaginella moellendorffii* (Semo), *Streptofilum capillatum* (Stca), *Volvox carteri* (Voca).

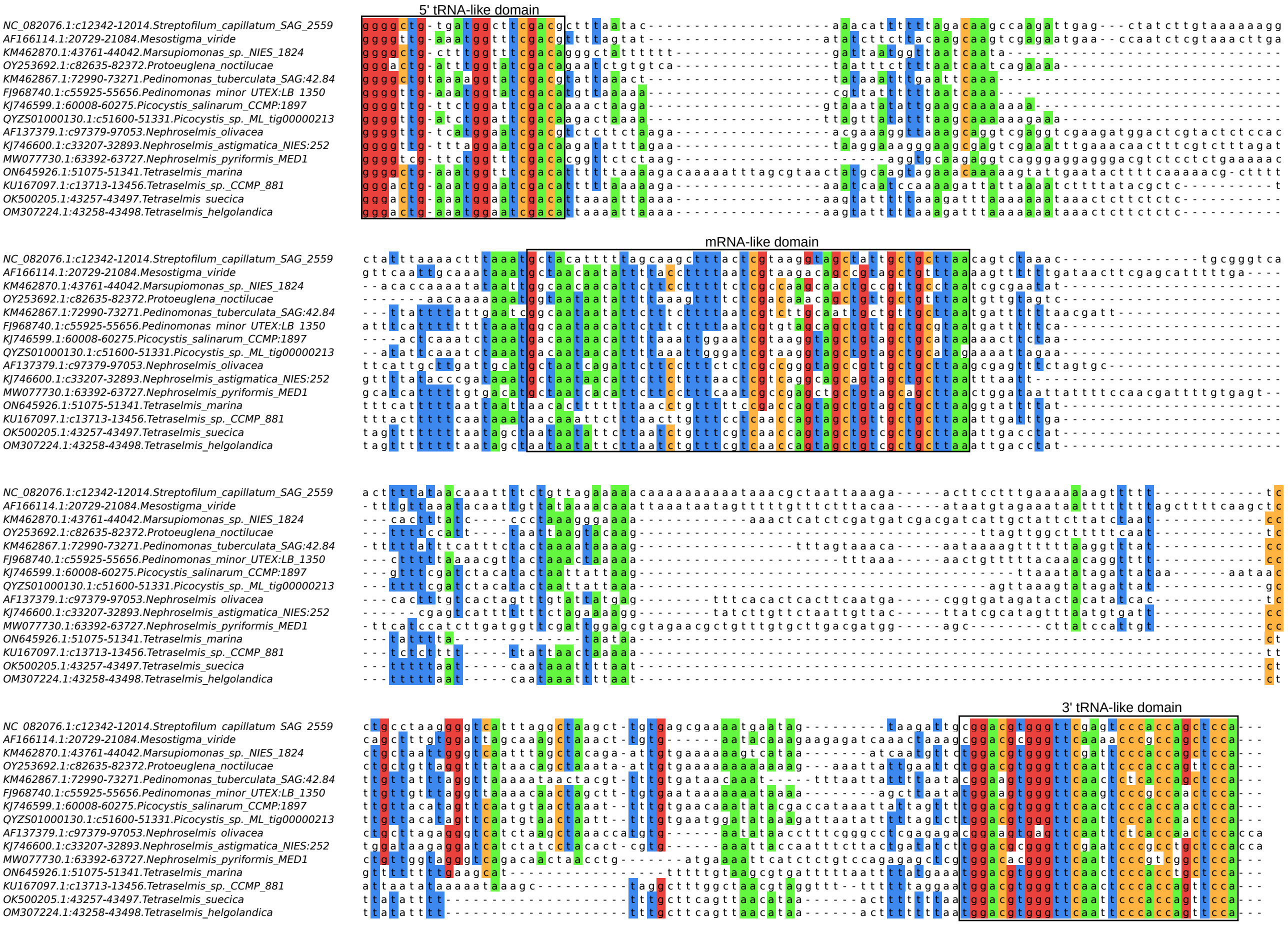


**Figure S4. The presence of the ssrA gene in the plastome of *Streptofilum capillatum* SAG 2559.** A previously unrecognised tmRNA-specifying gene (*ssrA*) occurs in the plastid genome of *S. capillatum* in the region flanked by the *ycf3* and *trnMe(cau)* genes. Presented is an alignment of the *S. capillatum ssrA* sequence with homologous sequences from other green algal plastomes, including *ssrA* genes identified for the first time here (those from *Tetraselmis* spp.). Accession numbers of the corresponding plastome sequences are provided, with the regions representing the *ssrA* gene specified by coordinates (the letter "c" means the gene is on the complementary strand). Boxed are regions corresponding to the conserved parts of the tmRNA molecule. The 5' and 3' regions pair to form the tRNA-like domain, which serves as a bona fide tRNA, bringing an alanine residue to the ribosome. To this end, an alanylatable 3' CCA end must be present in the molecule, which is presumably directly specified by the gene sequence in the case of *ssrA* from *Nephroselmis olivacea* and *Nephroselmis astigmatica*, whereas post-transcriptional extension of the tmRNA molecule by CCA tRNA nucleotidyltransferase must be assumed in the other cases. The mRNA-like domain in the middle is read by the ribosome upon resuming its translation activity, resulting in the addition of a short peptide tag to the nascent protein marking it for degradation. Note the presence of a termination codon (nearly always TAA) in the mRNA-like region.

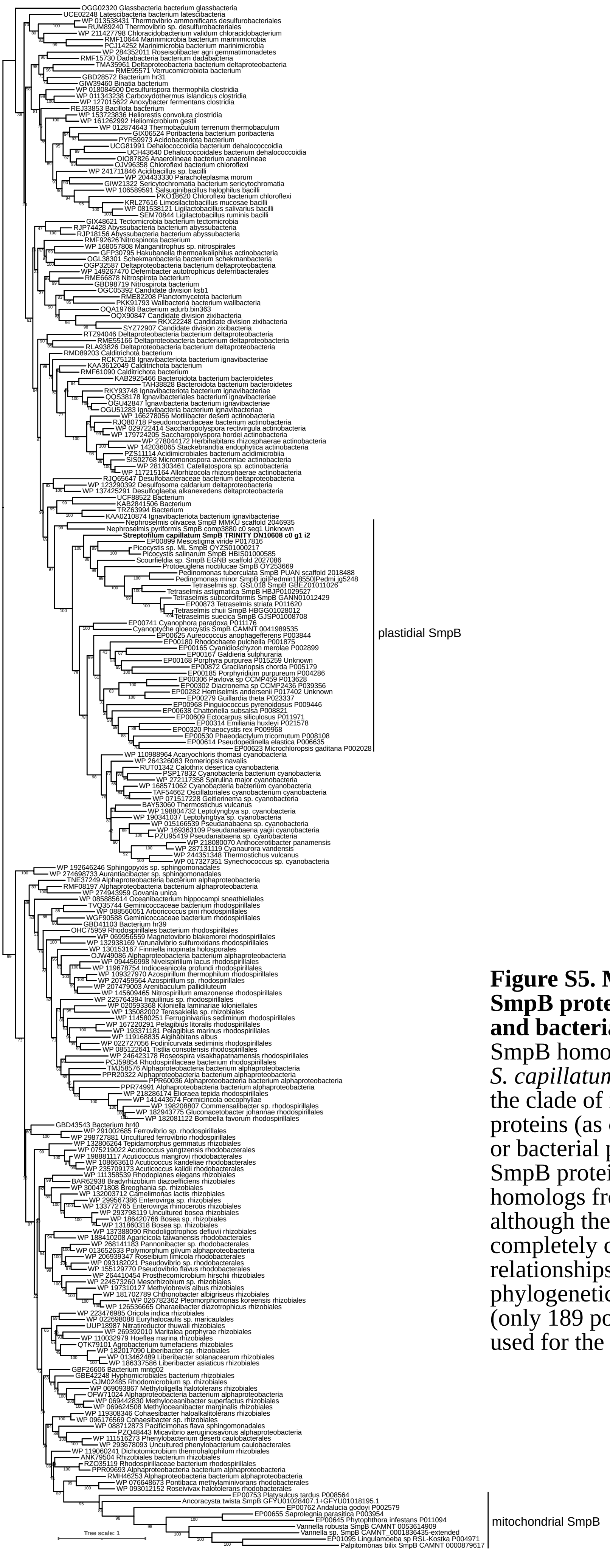


**Figure S5. Maximum likelihood phylogeny of SmpB proteins from eukaryotic organelles and bacteria.** The tree demonstrates that the SmpB homolog identified in the decontaminated *S. capillatum* transcriptome assembly belongs to the clade of nucleus-encoded plastid-targeted proteins (as opposed to mitochondrion-targeted or bacterial proteins). Note that the plastidial SmpB proteins are most closely related to homologs from Cyanobacteria, as expected, although the relationships among them is not completely congruent with the species relationships, presumably because of the limited phylogenetic signal in the short SmpB sequence (only 189 positions in the trimmed alignment used for the phylogenetic analysis).

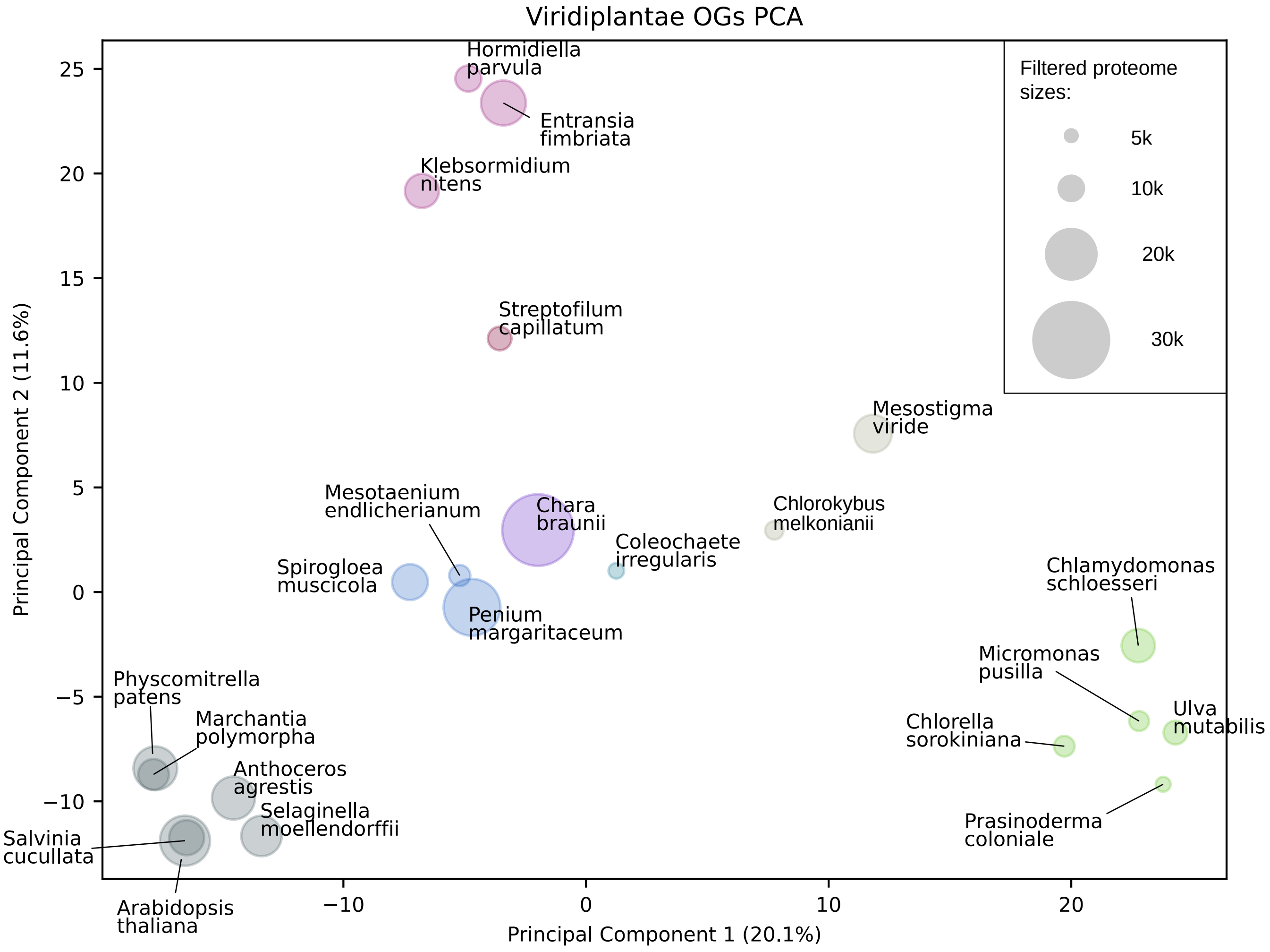


**Figure S6.** Comparison of protein repertoires (predicted proteomes) encoded by selected green algae and plants. This is a detailed version of a scheme displayed in Figure 1B (see the respective figure legend for further explanation).
